# Honey Bee *(Apis mellifera)* Exposomes and Biological Pathways Associated with *Nosema ceranae* Infection

**DOI:** 10.1101/152249

**Authors:** Robert L. Broadrup, Christopher Mayack, Sassicaia J. Schick, Elizabeth J. Eppley, Helen K. White, Anthony Macherone

## Abstract

A pilot study was conducted to determine if exposome profiles of honey bees *(Apis mellifera)* are associated with *Nosema ceranae* infection and whether xenobiotic exposures effect changes in known biological pathways of bees. Thirty stationary hives were selected from seven apiaries representing urban and suburban geographies. Foraging bees were harvested during the summer of 2015 and analyzed for *Nosema ceranae* infection via semi-quantitative polymerase chain reaction (sq-PCR) and discovery-based exposome analysis via gas chromatography-time of flight mass spectrometry (GC-TOF). The resulting datasets were divided into case and control groups based on the prevalence of *N. ceranae* infection. Xenobiotic burden was determined to be associated with *N. ceranae* infection, and co-variate analysis determined differentially expressed biological chemicals and naturally occurring chemicals in the bee exposomes. Biological pathways analyses putatively identified 10 dysregulated pathways as well as the presence of the P450 oxidative metabolism of naphthalene for detoxification. Based on these results, it is evident that the integration of genetic disease screening with discovery-based exposomics provides a promising multi-omic platform to identify adverse biological effects to bees occurring from exposures to chemicals and parasites. In addition, this approach will generate new hypotheses for targeted follow-up studies to examine bee health.

## Introduction

Honey bees are an essential part of global ecology, and their services as crop pollinators cannot be overstated^1^. Over the past decade global bee populations have been severely reduced by various environmental stressors including but not limited to: poor nutrition, losses in foraging habitats, infectious exposures to viruses and parasites and exposures to pesticides and other persistent chemicals^2, 3, 4^. Each of these individual stressors represents environmental exposures that adversely affect the health of bees at the colony level. Moreover, these environmental exposures may interact with one another, resulting in synergistic effects on the overall health of the hive^5, 6, 7^ A recent study indicated that the interaction between viral infections and the parasitic mite Varroa destructor, is a primary cause of honey bee colony collapse^8^. The probability of infection by the fungus, Nosema spp., has also been found to be substantially increased when bees are concomitantly exposed to fungicides^9^. Moreover, Nosema spp. infection coupled with exposure to the neonicotinoid pesticide imidacloprid has been observed to weaken the ability of honey bees to sterilize the colony and brood food, rendering the hive more susceptible to pathogens^10^. These studies suggest that there is a complex relationship between environmental exposures and hive survivability.

The exposome is the “life-course environmental exposures (including lifestyle factors), from the prenatal period onwards”^11^. It was espoused primarily to address the disproportionate characterization of the genome when compared to individuals’ environmental exposure data in cancer epidemiology. The exposome paradigm described the application of omics technologies for the characterization of non-genetic exposures to balance the gene (G) and environment (E) components of the Phenotype = G + E principle^12^. Five years after the exposome was first espoused, an influential perspective defined the “environment [with respect to the exposome] as the body’s internal chemical environment and exposures as the amounts of biologically active chemicals in this internal environment” and a “top-down strategy” to measure them^13^. In 2012, the definition of the exposome was stratified to include three domains: the general external (climate, financial status, education, etc.); the specific external exposome (pollution, diet, lifestyle factors including drug, alcohol abuse and tobacco use, etc.); and the internal exposome (metabolism, activity of the microbiome, oxidative stress, etc.)^14^.

Critical tenets of the exposome are to ascertain exposure-response relationships (biochemical epidemiology), mechanisms of action (systems biology), and the sources of exposure and kinetics (exposure biology)^15^. The exposome concept has been previously applied to the study of honey bee health using both targeted^16^ and untargeted^17^ approaches. Traynor, et al. (2016)^16^, used a targeted approach focused on pesticide exposures to study 91 colonies from three different migratory beekeeper operations that provide pollination services on the eastern coast of the United States. The bees were monitored longitudinally over the course of 10 months (March 2007 through January 2008). Samples of adult bees, beeswax, and beebread were collected and analyzed for 171 targeted pesticides. These data were used to determine associations between total pesticide burden and hive morbidity and mortality. They further explored the influence of these exposures on hive mortality and re-queening events by measuring the difference in the total number of pesticides at the beginning and at the end of the study. This work investigated the effects of the specific exposome domain as defined above (pesticide exposure) and, except for modes of action, no specific biological response (pathways) information was presented.

In this study, we examined honey bee exposome profiles from hives infected with *N. ceranae* and from uninfected hives. Statistical tests and bioinformatics tools available in commercial and open-source software were employed to determine associations between xenobiotic burden and *N. ceranae* load and, for pathways analyses.

## Results

### *Nosema ceranae* infection load

Eighteen of the thirty hives sampled for this study were found to have some level of *N. ceranae* infection but no *N. apis* was detected (see Supplemental Table S1). The *N. ceranae* results were log-normal distributed. The mean (± s.d.) was 3.99±9.22 the median value was 0.54, and the results ranged from 0.00 to a maximum of 40.02. Based on this data, the samples were divided into case (*N. ceranae* load > 0, n = 18) and control (*N. ceranae* load = 0, n = 12) groups.

### Xenobiotic burden

From the annotated chemical features, twenty known xenobiotics (chemicals not known to naturally occur in honey bees) were identified (Table 1). These twenty chemicals occurred a total of 143 times across all thirty samples. Each sample contained at least one xenobiotic, and the maximum number of xenobiotics identified in a single sample was eight. The average number of xenobiotics identified per sample was five. Figure 1 illustrates the cumulative number of exposures stratified by chemical category in the case and control groups.

**Table 1.** Xenobiotics identified in honey bee exposomes. Chemical category and CAS number provided for each.

| *Compound* | *Category* | *CAS Number* |
| --- | --- | --- |
| *Praziquantel* | antiparasitic | 55268-74-1 |
| *Captafol* | fungicide | 2425-06-1 |
| *Captan* | fungicide | 133-06-2 |
| *Fenpropimorph* | fungicide | 67564-91-4 |
| *Propamocarb* | fungicide | 25606-41-1 |
| *Carbetamide* | herbicide | 16118-49-3 |
| *Benzoylprop-Ethyl* | herbicide | 22212-55-1 |
| *Tebutam* | herbicide | 35256-85-0 |
| *Allethrin* | insecticide | 584-79-2 |
| *Azobenzene* | insecticide | 103-33-3 |
| *Carbofuran* | insecticide | 1563-66-2 |
| *Ethiofencarb* | insecticide | 29973-13-5 |
| *Flurecol-Butyl* | insecticide | 2314-09-2 |
| *Isoprocarb* | insecticide | 2631-40-5 |
| *Monocrotophos* | insecticide | 919-44-8 |
| *Promecarb* | insecticide | 2631-37-0 |
| *Propoxur* | insecticide | 114-26-1 |
| *Tetramethrin* | insecticide | 7696-12-0 |
| *Oxalic Acid* | varroacide | 6153-56-6 |
| *Naphthalene* | PAH | 91-20-3 |

**Figure 1.**
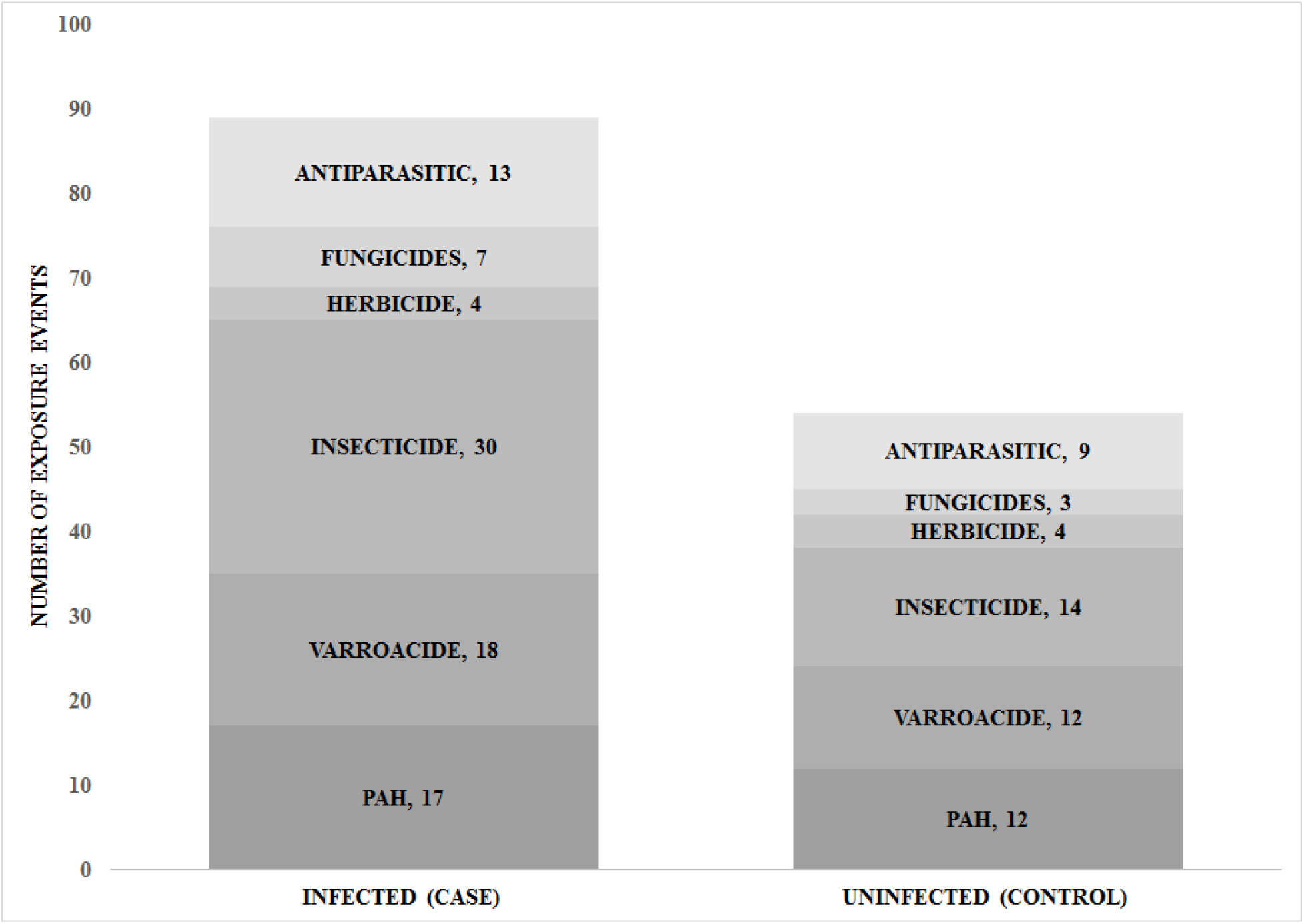
Cumulative number of xenobiotic exposure events. Xenobiotics identified in exposome profiles of honey bee foragers categorized by N. ceranae infected and uninfected hives. Infected hives are insulted with 65% more exposure events compared to uninfected hives. The various exposures represent 9 known modes of action.

### Xenobiotic burden and *N. ceranae* infection

#### Number of xenobiotic exposure events

In *N. ceranae* infected hives (case), 18/20 xenobiotics were found, and 10/20 were found in the uninfected hives (control). These data were used to determine that the total number of exposure events to individual xenobiotics varied significantly between the case and control groups (χ^2^ = 7.619, df = 1, n = 30, p < .006) (see Supplemental Table S2).

#### Relative levels of exposure in case and control groups

The data were further divided based on the number of total exposure events with a mass spectral ion abundance (representing a relative level of exposure for the case and control groups) greater than or less than the combined median ion abundance (24.36, log2 normalized). In this analysis, it was determined that the case and controls did not vary significantly (χ^2^ = 0.168, df = 1, n = 30, p =0.682) which suggests that the relative levels of exposure were not correlated with Nosema infection load (see Supplemental Table S3).

#### Relative levels of exposure in case and control groups stratified by chemical category

When the total number of exposure events with ion abundance greater than or less than the combined median ion abundance is stratified by chemical category, there was no correlation between the relative level of exposure and *Nosema* infection load (Table 2 and Supplemental Fig. S1 and Table S4).

**Table 2.**
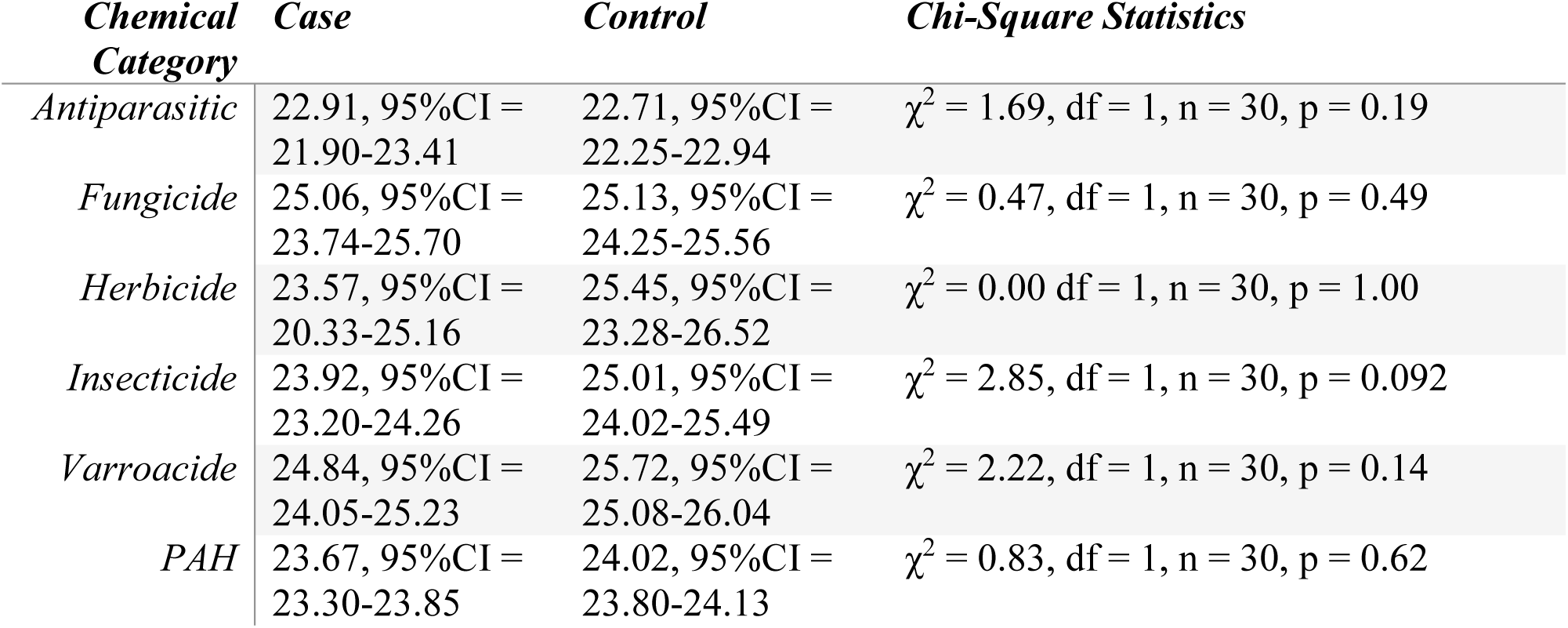
Chemical category and descriptive statistics. Average log_2_ normalized ion abundance with 95% confidence intervals for cases and controls and chi-square statistics.

### Putative identification of biological pathways relevant to honey bee health

Using the differentially expressed chemical entity sub-set of biological and naturally occurring chemicals in the bee exposome profiles determined through statistical testing and co-variate analysis, we identified 10 biological pathways and 14 biological chemicals associated with *N. ceranae* infection (Table 3). In addition, naphthalene, a known xenobiotic, was identified and associated with the P450 oxidation pathway.

**Table 3.** Pathways Analysis. Identified biological pathways and associated chemical entities.

| *Pathway* | *Chemical Entity Name* |
| --- | --- |
| *Biosynthesis of unsaturated fatty acids* | octadecanoic acid, (9Z)-octadecenoic acid, hexadecanoic acid |
| *Fatty acid biosynthesis* | octadecanoic acid, hexadecanoic acid |
| *Fatty acid degradation* | hexadecanoic acid |
| *Fatty acid elongation* | hexadecanoic acid |
| *Histidine metabolism* | N(pi)-methyl-L-hisidine, 4-(beta-acetylaminoethyl)imidazole |
| *Steroid biosynthesis* | beta-sitosterol, isofucosterol, squalene |
| *Sulfur Metabolism* | dimethyl sulfone, sulfur (cyclic octaatomic sulfur) |
| *Tyrosine metabolism* | 3,4-dihydroxyphenylethylene glycol |
| *Ubiquinone and other terpenoid-quinone biosynthesis* | beta-tocopherol |
| *Citrate (TCA) cycle* | succinate |
| *P450* | naphthalene |

### Modes of action

For the 20 xenobiotics identified in the bee exposomes, 9 modes of action (MOA) are known (see Supplemental Table S5). The MOA for the herbicide benzoylprop-ethyl and for Naphthalene (PAH) are unknown. Combining the 9 known MOA categories into 4 categories with a broader scope reveals a total of 112 xenobiotic exposure events, and infected hives were about 65% more exposed to xenobiotics compared to uninfected hives. The calculated p-values for a binomial distribution at α = .05, determined that the Ionic (Na^+^, Ca^2+^) Interference category was significantly associated (p < .01), and the Chemical / Enzyme Interference category may be associated (p < .05) with *Nosema* infection (Figure 2). The Multiple Modes of Action or Physiological Effects categories were not associated with *Nosema* infection, p = 0.12 and p = 0.23, respectively (Table 4).

**Figure 2.**
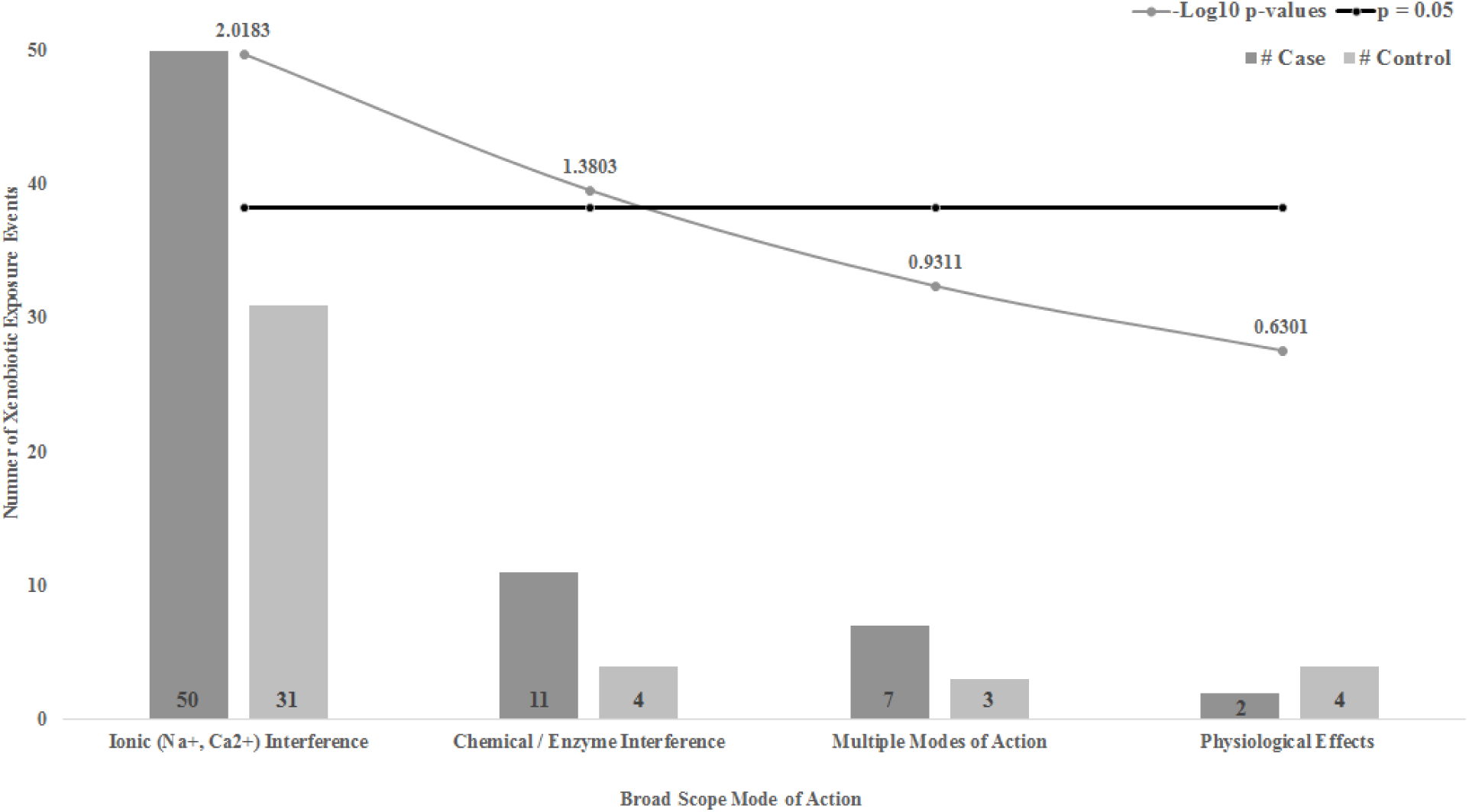
Broad categories of chemical exposures for case and control groups. The p-value for each category is given in the superimposed curve and the p = 0.05 cut-off is represented by the horizontal line. p-values given as -Log_10_ transformed.

**Table 4.** Broad MOA categories. The total number of exposure events (n) in each category, the number of exposure events stratified by case and control and, p-values and -Log_10_ p-values determined for each.

| ***Broad MOA***<br>***Categories*** | ***n*** | ***# Case*** | ***# Control*** | ***p*** | ***-Log<sub>10</sub> p*** |
| --- | --- | --- | --- | --- | --- |
| *Ionic (Na<sup>+</sup>, Ca<sup>2+</sup>)*<br>*Interference* | 81 | 50 | 31 | 0.0096 | 2.02 |
| *Chemical /*<br>*Enzyme*<br>*Interference* | 15 | 11 | 4 | 0.042 | 1.4 |
| *Multiple Modes*<br>*of Action* | 10 | 7 | 3 | 0.12 | 0.93 |
| *Physiological*<br>*Effects* | 6 | 2 | 4 | 0.23 | 0.63 |

## Discussion

It is generally accepted that pesticide exposures suppress the honey bee immune system, which increases the risk of contracting an infection^18, 19^. Xenobiotic exposures have various MOA that effect multiple genes and biological pathways along with effecting energy metabolism and cellular stress response. These physiological changes may increase susceptibility to other stressors such as *N. ceranae* infections that result in energetic dysregulation. The primary response to viral infections also includes changes in gene expression of metabolic pathways^20^, supporting the notion that dysregulation of metabolism is likely where the synergistic decline in bee health is occurring mechanistically.

We identified three differentially expressed chemicals (octadecanoic acid, (9Z)-octadecenoic acid, hexadecanoic acid; known components of beebread^21^) associated with fatty acid metabolic pathways (Table 3). Dysregulation of these pathways are known to play a role in immunocompetence^22^. Recently, it has been shown that adipokinetic hormone which is responsible for mobilizing fat stores is unresponsive in uninfected and infected honey bees, even under starved conditions^23^. These findings support the notion that honey bees may be particularly reliant upon a limited subset of metabolic pathways when responding to energetic stress, and disruptions of these pathways may have a substantial impact on health in *N. ceranae* infected bee colonies.

The lack of available energy in *N. ceranae* infected bees is one of the primary effects of chronic *Nosema* infection^24^, resulting in considerably lower trehalose levels in the hemolymph^25^. Interestingly, *N. ceranae* tolerant strains of honey bees can maintain stable trehalose levels in the hemolymph despite high parasite loads^26^. As *Nosema* infection is dependent on the host to supply ATP to reproduce^27^, it is not surprising that host metabolic pathways associated with carbohydrate metabolism are altered in response to infection. For example, changes in regulation of both gene expression and proteins such as alpha-glucosidase II, that are directly involved with carbohydrate catabolism and the generation of ATP through the electron transport chain, are consistently upregulated in previous studies^28, 29, 30^, and these changes may affect ubiquinone and terpene metabolism and differential expression of beta-tocopherol. We also observed differential expression of sulfur and dimethyl sulfone in this pathway. Dysregulation of sulfur metabolism may be due to exposures to sulfur-containing environmental pollutants.

Fungicides interfere with nutrient acquisition and metabolism; therefore, bees are likely to suffer from malnutrition even though adequate pollen may be available^31^. On their own, sub-LD50 (if known) exposures to fungicides are safer for bees in comparison to insecticide exposures, but they are known to have increased damaging effects when combined with other stressors^32^. Colonies exposed to low levels of fungicides often exhibit poor brood rearing, colony weakening, poor nutrition acquisition, and increased virus titers that together can lead to colony loss ^33, 34, 35^. Studies have shown ATP levels are impacted when bees are exposed to fungicides ^16, 36, 37^ and suggest that these changes are likely due to inhibition of succinate dehydrogenase^38^, which bridges the highly-conserved TCA cycle to the electron transport chain required for ATP generation. Malnutrition in bees is also associated with increased susceptibility to the effects of pesticides.

Tyrosine dysregulation may be a result of apiaries supplementing food sources with high fructose corn syrup or sucrose. Genes have been identified that are associated with xenobiotic detoxification and tyrosine metabolism^39^, and xenobiotics exposures have been shown to up-regulate proteins involved with detoxification processes, energy, carbohydrate, and amino acid metabolism^40^. We identified up-regulation of succinate in the TCA cycle that may be due to nutritional deficiencies or supplements. Two apiaries were known to supply sucrose feed but we have no information from the others about nutritional supplementation.

Relatively little is known about how multiple pesticide residue exposures impact the life stages of honey bees or wild pollinators. Previous studies examining the synergistic effects between parasites and pesticide exposure have been tested in the laboratory, but the results remain unclear as they have not been determined to exceed naturally occurring stochastic events^41^. For all foragers sampled in this work, we consistently detected older classes of pesticides such as carbamates, pyrethroids, and organophosphates. Additionally, the xenobiotics identified in our study differ somewhat from those reported in previous studies^42^. For example, we did not detect coumaphos, a common insecticide with known acetylcholinesterase inhibition activity, in the foraging bees. This may reflect our sample set being comprised of hives from beekeepers located in suburban and urban areas as opposed to commercial hives that are often transported for pollination services in more rural locations.

We identified a total of 44 insecticide exposures (30 case, 14 control) and only 10 fungicide exposures (7 case, 3 control) in our dataset. These findings agree with a recent study that demonstrated how bees sampled from the field are more likely to be exposed to insecticides encountered through diverse foraging of wildflowers and other pollen sources (non-focal crop pollen) than fungicides which dominate focal crop pollen foraging^43^.

Both xenobiotic exposures and exposures to parasites such as *N. ceranae* are considered part of the measurable specific external exposome. Even though both *Nosema* infected and uninfected hives were exposed to the same xenobiotic chemical space and both groups contained xenobiotics representing all 6 chemical categories, our data suggest that xenobiotic burden associated with *N. ceranae* infection may depend more on the number of exposures than the actual exposure type (chemical category). Our data further suggest that the relative level of exposure as estimated by the ion abundance in the mass spectra of the bee extracts is poorly correlated to *Nosema* infection. However, we cannot rule out the cumulative effect of a high number of exposure events in the case samples that represent many modes of action effecting bee physiology and possibly overwhelming the bees’ ability to recover from multiple stressors – thus leading to infection.

This study demonstrates how xenobiotic exposures may act synergistically to increase susceptibility to disease, and serves as a proof of concept for the integration of genetic disease screening with discovery-based exposomics using TOF mass spectrometry. Through a multi-omic integration of data, we can observe how xenobiotic exposures effect bee physiology through multiple modes of action rendering the bees more susceptible to environmental pathogens such as *Nosema ceranae*. This analytical platform measures and characterizes the effects of exposures from the specific external exposome (e.g., xenobiotics and *N. ceranae*) in the internal exposome environment of the bees (bee extracts). These data identify changes in the metabolome and provide a conduit for the identification of affected biological pathways. We postulate the changes in the internal environment of the bees results in a negative chemical feedback loop (Figure 3) and further increase susceptibility to infection and ultimately, the decline of hive health.

**Figure 3.**
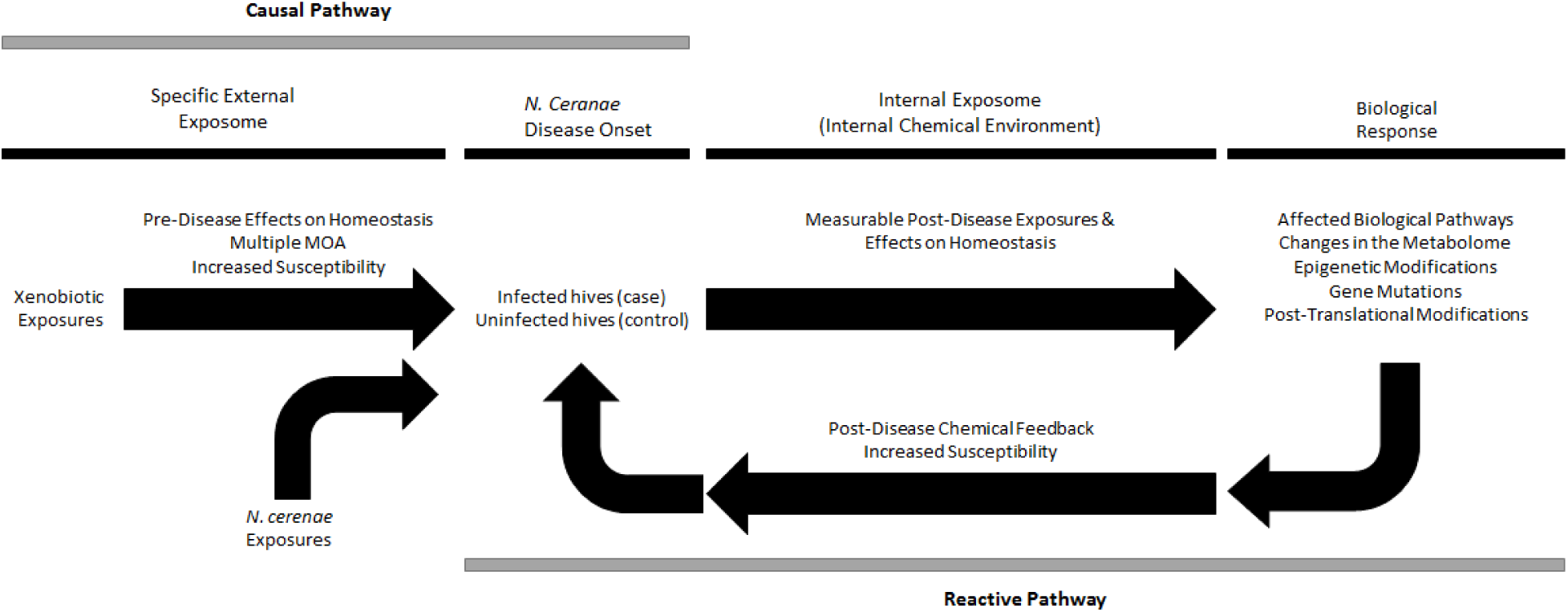
Disease pathways^56^. Along the pre-disease causal pathway, specific external exposures effect measurable chemical changes and increase susceptibility to disease through multiple MOA. These exposures affect biological pathways, change the metabolome, and influence genetic expression, transcription, and post-translational and epigenetic modifications. Along the post-disease reactive pathway, measurable chemical changes in the various “omes” feedback and further increase disease susceptibility in an increasingly adverse feedback loop.

## Methods

### Sample Collection

Adult foraging bees were sampled at a single time point during the 2015 season from 30 hives in 7 different geographical locations in the southeastern Pennsylvania region. The locations were selected to represent urban and suburban settings (see Supplemental Table S6). Sixty to one hundred bees in total were collected from each hive in 50 mL disposable tubes and immediately frozen on dry ice. All samples were then stored in a -80° C freezer. Prior to analysis the samples from each hive were gently thawed and divided into two groups of 30 bees: one group for sq-PCR analyses and a second group for extraction and sample cleanup prior to GC-TOF analysis. Each of the resulting 60 samples (30 hives x 2 groups/hive) represent a snapshot of the parasitic load and exposome profile of the hive from which it was collected at that specific time. All samples were randomized prior to analysis.

### Honey bee Extraction for GC-TOF analysis

Honey bees were extracted using previously published QuEChERS extraction protocols^44, 45^. Briefly, 3 grams of foraging bees (~30 bees) were pulverized in 27 mL 44:55:1 water/acetonitrile/acetic acid and transferred to 50 mL disposable tubes. Six grams of magnesium sulfate and 1.5 g sodium acetate were added to the suspension, and the tubes were sealed, shaken, and thoroughly mixed before centrifugation at 3000 rpm for 5 minutes. A 2-mL aliquot of the supernatant from each sample was applied to conditioned C18 SPE cartridges (Agilent Technologies, Santa Clara, CA). The analytes were eluted with a 70:30 solution of acetone/toluene and reduced in volume before transfer to 2 mL auto-sampler vials prior to GC-TOF analysis.

### GC-TOF

For discovery-based (non-targeted) exposome profiling of honey bee extracts, an Agilent 7890B/7200B gas chromatography-quadrupole time of flight mass spectrometer (GC-QTOF) system was used. A 0.2 μL pulsed split-less injection was made into a 250 °C isothermal split/split-less inlet. The GC was configured with a 40 m x 0.25 mm x 0.25 μm DB5-MS DuraGuard column (J&W 122-5532G) operated at 1.2 mL/minute helium in constant flow mode. The oven program was 80 °C (1 minute) then, 10 °C/min to 310 °C (6 minutes). The transfer line temperature was 300 °C. The mass spectrometer was operated in electron ionization, high resolution TOF mode. The source and quadrupole (RF only) temperatures were 275 °C and 150 °C, respectively. High resolution, accurate mass (HRAM) spectral data was collected at 5 Hz over a mass range of 50 Da to 800 Da. Automated intra-sequence mass calibration was performed immediately prior to each sample injection. All samples were run in duplicate and the average of the two runs used for data analysis.

### Chromatographic deconvolution and chemical entity annotation

Raw data acquired on the GC-QTOF system was analyzed using the MassHunter suite of software. To this end, Unknowns Analysis B.08.00 was used to perform chromatographic deconvolution of the 60 data files collected in this pilot study (30 samples x 2 injections per sample). Chemical features were minimally identified as having signal-to-noise ratio > 3:1, an accurate mass assignment for the base ion and a retention time for the chromatographic peak where the feature is found. These criteria identified a total of 2,352 chemical features.

Spectral library searches and compound annotation were performed using the RTL Pesticides and the Fiehn Metabolomics libraries (Agilent Technologies, Santa Clara, CA) and the NIST-11 Mass Spectral Library (the National Institute of Standards and Technology, NIST Standard Reference Database 1A v11). Of the 2,352 identified chemical features, 1,723 (73%) were annotated (retention time, ion abundance, m/z, chemical name, CAS number) by spectral library search. For the 629 chemical features that were not identified, the minimal feature parameters defined above and a composite mass spectrum was used for covariate statistical analysis.

### Statistical testing and covariate analysis

Identifying biological associations with the myriad of exposures encountered over individuals’ lifetime in a hypothesis-free (data-driven) manner requires sophisticated bioinformatics tools. It has been demonstrated how data-driven discovery of disease-exposure associations can generate new hypotheses through Environment-Wide Association Studies (EWAS),^46^ which have also been referred to as Exposome-Wide Association Studies^47^.

To identify associations of exposome profiles with *N. ceranae* infection, the mass spectrometry datasets collected in this study were statistically analyzed with MassProfiler Professional (MPP) bioinformatics software to identify chemical features associated with the *N. ceranae* infected samples. This process entailed data alignment, baselining, quality testing and significance testing (unpaired T-test or one-way ANOVA, p < .05). Only chemicals that passed the above filters and statistical testing with a relative ion abundance fold-change at least two-times the median ion abundance across all samples remained in the data subset. These associated chemicals were then screened against known *Apis mellifera* biological pathways obtained from the Kyoto Encyclopedia of Genes and Genomes (KEGG) database ^48, 49^. Other data analyses and graphics generation were performed using Microsoft Excel^®^ 2016, R and R Studio software^50, 51^.

### Semi-quantitative-PCR screening for Nosema spp

#### DNA Extraction

Pooled samples for semi-quantitative PCR analyses were prepared by adding 6 mL of RNase free water to 30 bees collected from each hive. The bees were thoroughly homogenized with a sterile 50 mL tissue grinder (Fisher). Specific primers that identify and distinguish *N. apis* and *N. ceranae* based on a unique sequence found in a highly conserved ribosomal gene were used^52^. To semi-quantify both *Nosema ceranae* and *N. apis*, a PCR multiplex reaction with a RpS5 reference gene (see Supplemental Table S7) was performed based on methods adapted from the HBRC method^53^. Briefly, a 150 μL aliquot of the homogenate was added to 300 μL of a 1:1 mixture of phenol/chloroform. The solution was centrifuged at 13,000 rpm for 5 min and the supernatant was added to another 300 μL of the 1:1 phenol/chloroform solution in a new 1.5 mL microcentrifuge tube. The supernatant was drawn once again and transferred to a new 1.5 mL microcentrifuge tube that was then followed by the addition of 30 μL sodium acetate and 600 μL 95% ethanol to precipitate the DNA overnight at - 20 °C. For the PCR reaction, the extracted DNA from each sample was quantified using a Nanodrop 2000 UV-Vis Spectrophotometer (Thermo Scientific). Each sample was then diluted with DNase-free water to a working concentration of 5 ng/μl to serve as the PCR template DNA.

#### Semi-quantitative-PCR

A 15 μL duplex PCR reaction (one for *N. ceranae* and one for *N. apis*) combined: 1 μL of a 10 mM solution of each of the four primers (4 μL total volume); 1.5 μL of 10x PCR buffer; 0.5 μL of 10 mM deoxynucleotide triphosphate (dNTP); 0.2 μl of a 25 mM magnesium chloride solution; 0.2 μL of 5 U/μL Taq^®^ DNA polymerase (New England BioLabs, Ipswich, MA) with 2 μL of template DNA from the DNA extraction described above and 6.6 μL Millipore water (0.2 μm sterile filter).The PCR thermocycler program was 94 °C for 2.5 min, followed by 10 cycles of 15s at 94 °C, 30s at 61.8 °C and 45s at 72 °C, and 20 cycles of 15s at 94 °C, 30s at 61.8 °C and 50s at 72 °C, an extension step at 72 °C for 7 min, and a final hold step of 4 °C. Each PCR assay included a negative and positive control for each target.

PCR products were confirmed by a 3% gel electrophoresis at 100 v for 3.5 hours with a low molecular weight DNA ladder (New England BioLabs, Ipswich, MA). Each sample was run in triplicate and the relative amount of DNA was determined using densitometry and Image J software^54, 55^. Values were averaged across the triplicates and divided by the *Apis mellifera* RpS5 reference gene to calculate the relative abundance of *N. ceranae* or *N. apis* for each hive sampled.

#### Modes of action

We combined the 9 known MOA categories associated with the 20 identified xenobiotics into 4 categories with a broader scope, i.e. MOA affecting Na^+^ or Ca^2+^ were grouped into an Ionic (Na^+^, Ca^2+^) Interference category, MOA affecting mitosis or microtubule assembly were grouped into a Physiological Effects category, MOA affecting acetylcholinesterase inhibition, oxidative phosphorylation inhibition, or interference with ATP production were grouped into a Chemical / Enzyme Interference category and lastly, Multiple Modes of Action remained unchanged. We determined the number of exposure events for the case and control groups for each of the broad MOA categories. Presuming the probability of xenobiotic exposures is binomially distributed in each of the 4 broad categories, i.e. there is an equal probability that an exposure event in each category will affect the onset of *Nosema* or it will not, we calculated p-values for α = .05 to assess the association between the number of exposures in each category.

## Data Availability

The datasets generated during the current study are available from the corresponding author on reasonable request. All data analyses determined during this study are included in this published article (and its Supplementary Information files).

## Author Contributions

R. L. B. performed all field work, honey bee extractions, GC-TOF data collection, and contributed to the manuscript. C. M. performed PCR assays, biometric analyses and contributed to the manuscript. S. S. assisted with PCR assays and contributed to the manuscript. E. E. assisted with PCR assays. H. K. W. contributed to and edited the manuscript. A. M. designed the research idea, performed GC-TOF analysis, analyzed and evaluated the data and contributed to the manuscript.

## Acknowledgements

We would like to thank Chloe Wang, Malia Wenny, Alexis Schafsnitz, Naomi Chaqueco, and Katiana Rufino of Haverford College for their assistance in the field and the laboratory.

## Conflict of Interest

Authors declare no conflict of interest.

